# An interpretable and versatile machine learning approach for oocyte phenotyping

**DOI:** 10.1101/2022.02.01.478709

**Authors:** Gaelle Letort, Adrien Eichmuller, Christelle Da Silva, Elvira Nikalayevich, Elsa Labrune, Jean-Philippe Wolf, Marie-Emilie Terret, Marie-Hélène Verlhac

## Abstract

Meiotic maturation is a crucial step of oocyte development allowing its potential fertilization and embryo development. Elucidating this process is important both for fundamental research and assisted reproductive technology. However, only few computational tools, based on non-invasive measurements, are currently available to characterize oocyte meiotic maturation. Here, we develop a computational framework to phenotype oocytes based on images or movies acquired exclusively in transmitted light. We first trained neural networks to segment the contours of oocytes and their zona pellucida using a diverse cohort of both mouse and human oocytes. We then defined a comprehensive set of morphological features to describe a single oocyte. We have implemented these steps in a versatile and user-friendly open source Fiji plugin available to the mouse and human oocyte community. Then, we present a machine learning pipeline based on selected features to automatically recognize oocyte populations and determine their morphological differences. Its first application is a novel approach to screen oocyte strains and automatically identify their morphological characteristics. We demonstrate its potential by phenotyping a well characterized genetically modified mouse oocyte strain. Its second application is to predict and characterize the maturation potential of oocytes. Importantly, we identify two new features to assess mouse oocyte maturation potential, consisting in the texture of the zona pellucida and the cytoplasmic particles size. Eventually, we tested whether these mouse oocyte quality features were applicable to human oocyte’s developmental potential.

## Introduction

At the end of its growth in the ovary, an oocyte follows a critical phase called meiotic maturation which determines its capacity to be fertilized and sustain early embryonic development. Meiotic maturation consists of two successive highly asymmetric divisions in size (Mogessie et al., 2018; Verlhac and Terret, 2016). The first meiotic division (meiosis I) begins for an oocyte initially in prophase I with the rupture of the nuclear envelope (Nuclear Envelope BreakDown, NEBD, Figure 1A) and finishes with the formation of two daughter cells: a large oocyte and a small polar body (PB) (PB extrusion, Figure 1A). This highly asymmetric division allows the oocyte to retain the vast majority of its cytoplasmic content accumulated during its growth, essential for early embryonic development. The oocyte then enters into the second meiotic division (meiosis II) and arrests in metaphase II until fertilization by the sperm. This maturation step is particularly error-prone in terms of chromosome segregation and responsible for most aneuploidies in human (Mihajlović and FitzHarris, 2018; Nagaoka et al., 2012). Strikingly, in humans the quality of oocytes globally decreases with maternal age, which constitutes a major societal issue in modern societies where women tend to post-pone child bearing. It is thus the object of intense research efforts both in clinics and in academia, where it often relies on using surrogate models, close to human, to quantitatively analyze genetically modified oocytes.

**Figure 1:**
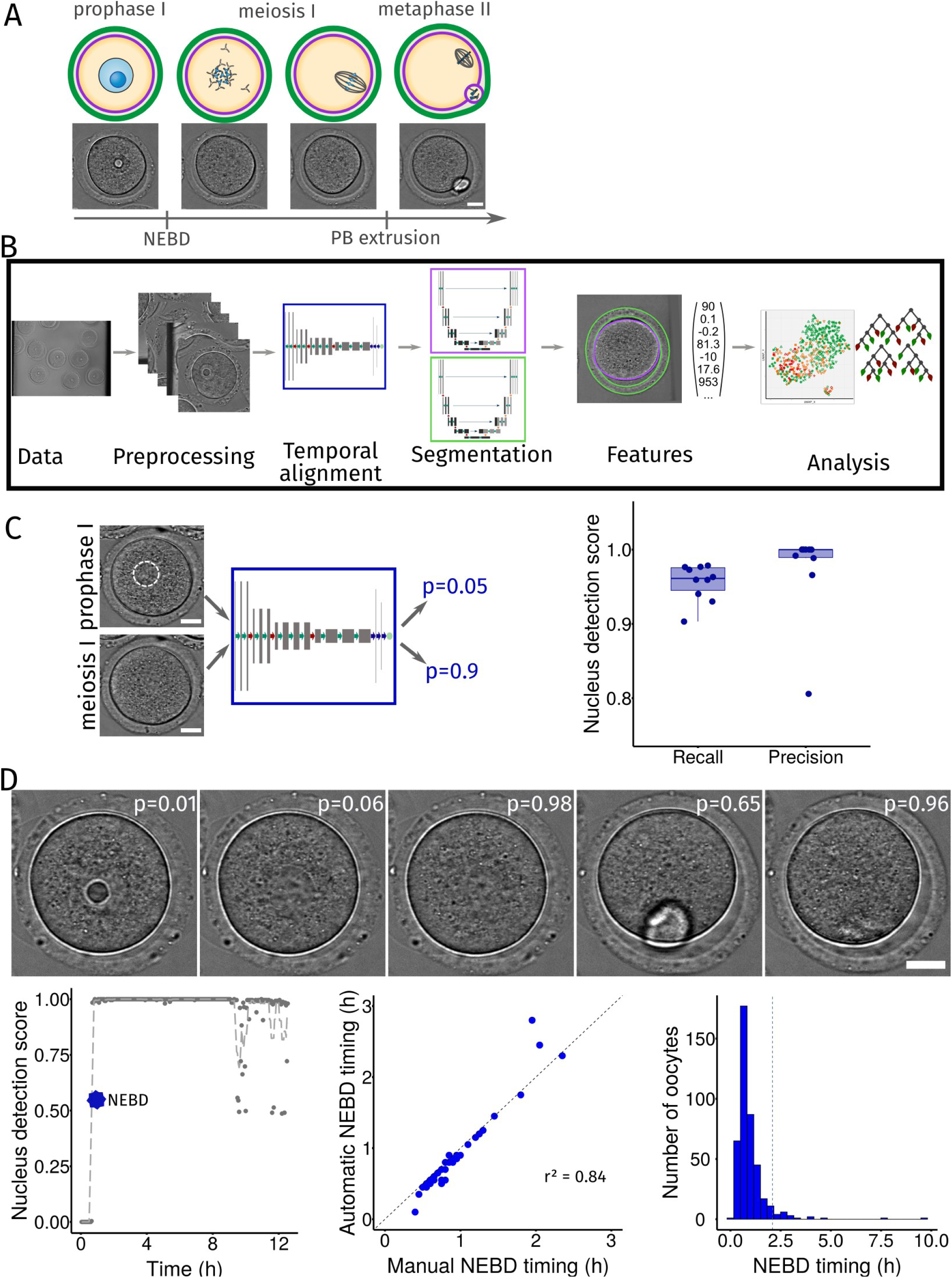
A machine learning pipeline to characterize oocyte developmental potential. A – Scheme of the main steps of oocyte maturation, from prophase I to metaphase II. Schematic representation (top) and still images from a movie of a mouse maturing oocyte in transmitted light (bottom). Scale bar is 20 µm. NEBD : nuclear envelope breakdown. PB: polar body. Blue: DNA, dark gray: microtubules, purple: oocyte plasma membrane, green: zona pellucida. B – Scheme of the machine learning pipeline. Steps used in our pipeline: Preprocessing, Temporal alignment, Segmentation, Features and Analysis. C – Automatic nucleus detection from transmitted light images. Training of the neural network (blue box, left panel) to recognize oocytes with a visible nucleus (upper left, the nucleus is indicated by the white dotted circle, bottom left, the oocyte does not have a nucleus). Scale bar is 20 µm. Scores (right graph) of nucleus detection on the test dataset (Recall: TP/(TP+FN) and Precision: TP/(TP+FP), where TP=True Positive, FN=False Negative, FP=False Positive) of 10 neural networks after training. D – Automatic determination of the NEBD timing in mouse oocyte maturation movies. Nucleus detection score (p, top panel) in individual images of one movie calculated with the trained neural network. Scale bar is 20 µm. Evolution of the nucleus detection score in time and automatic determination of NEBD timing as the first transition from low score (before NEBD) to high score (after NEBD, bottom left panel). Comparison of automatically and manually determined NEBD timing on 46 test oocyte maturation movies (bottom middle panel). R-square of the timing differences is indicated. The dashed line represents the y=x line. Histogram of the timing of NEBD from 424 movies (bottom right panel). The dashed vertical line represents the 0.95 quantile above which oocytes are considered to present a delay in NEBD timing.

In the context of assisted reproductive technologies, *in vitro* oocyte maturation can be used to avoid hormonal treatments for women experiencing repeated *in vitro* fertilization failures or who do not respond to hormonal stimulation (Hardy et al., 2000; La et al., 2019; Lonergan and Fair, 2016; Hatirnaz et al., 2018). Moreover, oocytes can also be extracted and frozen for fertility preservation. The success rate of oocyte *in vitro* maturation is typically around 70% (Kim et al., 2004), but the developmental potential of *in vitro* matured oocytes is much lower than for *in vivo* matured ones (Kim et al., 2004; La et al., 2019; Monti et al., 2017). To improve *in vitro* oocyte maturation protocols, we need to better decipher the parameters associated with successful oocyte development. Several characteristics correlating with oocyte quality have been identified, such as the composition and amount of accumulated maternal mRNAs, the size of the oocyte and its zona pellucida, its cytoplasmic organization, or the presence of certain epigenetic modifications (Cran, 1985; Eppig et al., 1994; Fair et al., 1995; Hyttel et al., 1989; La et al., 2019). However, a consensus on the individual power of these parameters to predict oocyte developmental potential has never been reached, and often with conflicting results (Rienzi et al., 2011). Machine learning techniques can be used to tackle this problem since they can automatically identify more relevant features and take into account multiple features at the same time. These techniques have been applied to predict metaphase II oocyte or blastocyst quality and select the best embryo for implantation (Khosravi et al., 2019; Wang et al., 2019; Cavalera et al., 2019; Manna et al., 2013), but not to predict prophase I oocytes maturation potential (Fernandez et al., 2020; Zaninovic and Rosenwaks, 2020).

In both academia and in clinics, computational tools to better characterize oocyte’s developmental potential are thus necessary. To follow live oocyte development, fluorescence imaging is commonly used in academia. However, it is limited to the number of fluorescent markers that can be followed in parallel and it produces phototoxicity potentially affecting oocyte’s proper development (Velilla et al., 2002; Kasprowicz et al., 2017; Ounkomol et al., 2018; Skylaki et al., 2016; Vicar et al., 2019). Label-free imaging is a promising non-invasive method to overcome these hurdles (Kasprowicz et al., 2017) and is exclusively used in clinics. However, label-free images present a much lower contrast, making the segmentation of the object contour a limiting step (Buggenthin et al., 2013; Tse et al., 2009). Deep learning approaches provide a good performance for segmentation of label-free images from various cell types (Falk et al., 2019; Kim et al., 2019; Ronneberger et al., 2015). Therefore, we chose a versatile approach to implement a segmentation tool robust to the diversity of oocyte types and modes of acquisition (mouse oocytes coming from academia and human oocytes from clinics), as well as to external objects present in the field of view such as follicular cells or injection pipettes.

Here, we developed a machine learning approach to characterize and predict oocyte maturation *in vitro* based on images acquired non-invasively. First, we propose a new Fiji plugin, Oocytor, available as open-source on github (see Methods). Oocytor is a user-friendly tool to segment the contour of an oocyte and of its zona pellucida from transmitted light images, to extract numerous morphological features and to automatically detect the rupture of the nuclear envelope marking the beginning of the maturation process. Next, we developed a machine learning approach to characterize and predict oocyte development. We decided to favor interpretability over predictive power and chose a machine learning approach based on interpretable features extracted from our plugin. We designed a list of 118 morphological features identified as potential markers of oocyte quality (Ozturk, 2020; Rienzi et al., 2011). We showed that this machine learning approach can be used to discriminate oocyte populations and automatically identify morphological differences coming from a mutant strain. This automatic phenotyping could be a valuable tool for fundamental research studies. Finally, we used our pipeline to predict the potential of oocytes to start and perform a correct maturation. In addition to predicting oocyte quality before its maturation, it allowed to identify the most determinant morphological features controlling the maturation process. As a proof of concept, we based our study on *in vitro* maturation of mouse oocytes, relatively close to human (Anderiesz et al., 2000; Levi et al., 2013; Ménézo and Hérubel, 2002) and offering access to more data and possible genetic manipulations. We tested how the results obtained on mouse oocytes could be transferred to *in vitro* human oocytes with a small dataset acquired from clinics.

## Results

To maximally avoid perturbations and to be in line with clinical practice, our method to analyze oocyte development was based on non-invasive measurements. Therefore, we propose here a computational tool to characterize oocytes from images or movies acquired in transmitted light, without any fluorescent marker. Our first objective was to build a user-friendly computational tool to extract quantitative information from images/movies of oocytes undergoing maturation. For this, we acquired 468 movies of mouse oocytes, starting shortly before nuclear envelope breakdown until metaphase II onset (Figure 1A).

### Overview of our new machine learning pipeline to characterize oocytes

To automatically analyze oocyte maturation, we propose a machine learning pipeline (Figure 1B overview of the pipeline and its different steps) based on our image data set. First, movies/images were preprocessed for homogenization (Material and Methods). Then, the time of maturation initiation, characterized by NEBD, was automatically determined and used as a temporal landmark (Figure 1B, Temporal alignment step). Next, images were segmented to identify the oocyte and its zona pellucida contours (Figure 1B, Segmentation step), allowing the extraction of hundreds of numerical features describing an oocyte (Figure 1B, Features step). We then used these features, or an uncorrelated subset of them, in several machine learning methods according to our needs (Figure 1B, Analysis step). We implemented the steps of Temporal alignment, Segmentation and Features into a Fiji plugin, Oocytor, proposing a user-friendly source to extract quantitative information from oocytes in transmitted light. Below we present more details on these 3 main steps of our pipeline.

### Automatic detection of NEBD from mouse oocytes

Oocyte maturation is a precisely temporally regulated process. The time after NEBD is commonly used as a landmark to describe progression into oocyte maturation. When maturation is triggered *in vitro*, the oocyte population does not undergo NEBD perfectly synchronously, with few oocytes extremely delayed and some remaining arrested in prophase I. Timelapse movies of oocyte development were temporally aligned on the NEBD timing to allow a quantitative temporal analysis of meiotic maturation progression. We therefore designed an automatic assessment of NEBD in our plugin to annotate this event.

We first trained a neural network to determine the presence or absence of a visible nucleus. Our network takes as input an image containing one oocyte and evaluates the probability that it has already undergone NEBD (Figure 1C, left panel, nucleus highlighted in dotted white circle). The architecture of a neural network determines its performance for a given task. We chose an architecture following VGG-16 (Simonyan and Zisserman, 2015), specialized for image classification (Figure S1A). We generated a dataset of mouse oocyte images manually annotated as before (0) or after (1) NEBD from our dataset (Material and Methods). Our final database comprised 7713 images, 90 % of which were used for selection and training of the network (Figure S1B, Material and Methods) and 10 % for testing its performance. We measured network precision and recall (Taha and Hanbury 2015) which give an indication of the quality and quantity of “hits” (here, oocytes after NEBD). We obtained a higher precision than recall (median score of 98% vs 96%, Figure 1C right panel), indicating that the network had a slight tendency to generate false negatives: oocytes without a nucleus predicted as still containing one. These false negatives might be due to the presence of a visible polar body in some images, which somehow confused the network. Indeed, the 4^th^ image of Figure 1D with a visible polar body has a lower score compared to the 3^rd^ and 5^th^ images without a visible polar body.

We corrected this problem by considering the overall information of the movie instead of looking only at a single snapshot and implemented NEBD determination in Oocytor. Each image of a movie was run through our trained neural network, which calculated a probability of being after NEBD for each time point (Figure 1D top panel). NEBD was set as the first transition point at which the score switches from low to high (Figure 1D bottom left panel). To evaluate the accuracy of our method, we compared the NEBD timing obtained with our plugin to the annotated one from test movies, which were not used for training the network. Overall, automatic and manual timings were very close (Figure 1D, bottom middle panel), which confirmed that our plugin can be reliably used for automatic detection of NEBD. Finally, we calculated the NEBD timing on our 468 mouse oocyte maturation movies. We manually checked the movies for which the obtained NEBD time was delayed. We found that 8 of the 44 oocytes that did not do NEBD were erroneously labeled as late NEBD and thus manually corrected. In the vast majority of the cases, oocytes ruptured their nuclear envelope (424 cells) approximately 30 min after the start of the recordings (Figure 1D, bottom right panel). However, some oocytes required more time to break their nuclear envelope, which might reflect states of unfinished growth or other defects. Taking the delay in NEBD timing into account is important since it can reflect inherent differences in oocyte potential and it is also essential to make comparison between oocytes at the same stage of meiotic maturation. Oocytor allows to do it automatically and avoid manual annotation, which could be tedious for large datasets.

### A robust and generic segmentation pipeline of oocyte contours

The second step of our machine learning pipeline was to extract quantitative and interpretable information from images to better describe oocytes. For this, we first implemented a deep learning-based tool to segment the contour of the oocyte from images acquired in transmitted light. We based our neural network (Figure S2A) on the U-Net architecture (Ronneberger et al., 2015), specifically designed for segmentation of biomedical images (Caicedo et al., 2019; Saleh et al., 2019). First, we created a database of thousands of images of single oocytes with their corresponding segmentation (ground-truth). With the aim of building a robust and versatile network (Möckl et al., 2020), we pooled together datasets of both mouse and human oocytes, coming from different contexts (Material and Methods). We obtained a database of 8256 images from which 85 % were used for selection and training of the network and the remaining 15 % for testing it (Figure 2A). During training, the network learned by optimizing its parameters to minimize the error between its outputs and the ground truth images (Figure 2A, right panel). We evaluated this error by the intersection over union (IOU) score, a metric measuring segmentation quality (Taha and Hanbury, 2015). The implementation of the network requires to choose additional parameters (e.g. the number of iterations) called hyper-parameters. Their fine tuning was performed on the training dataset with a cross-validation technique (Material and Methods, Figure S2B). We then trained the selected network (Figure 2A left). For mouse oocytes, we obtained an average IOU of 97% on the test dataset (Figure S2D). We applied the same steps to select and train neural networks to determine the contours (inner and outer limits) of the zona pellucida, a glycoprotein layer surrounding the oocyte and obtained an average IOU of 82 % on the mouse test dataset (Figure S2C-D). Eventually, we implemented this step in our Fiji plugin with the objective of proposing a user-friendly tool for segmentation of oocyte and zona pellucida contours from transmitted light images to the oocyte community (Material and Methods, Figure S2E). Figure 2B shows examples of segmentation obtained with Oocytor for oocytes at different stages of maturation. Oocytor can thus be used to segment oocytes from transmitted light images and based on how it was constructed, it should be relatively robust to different imaging conditions.

**Figure 2:**
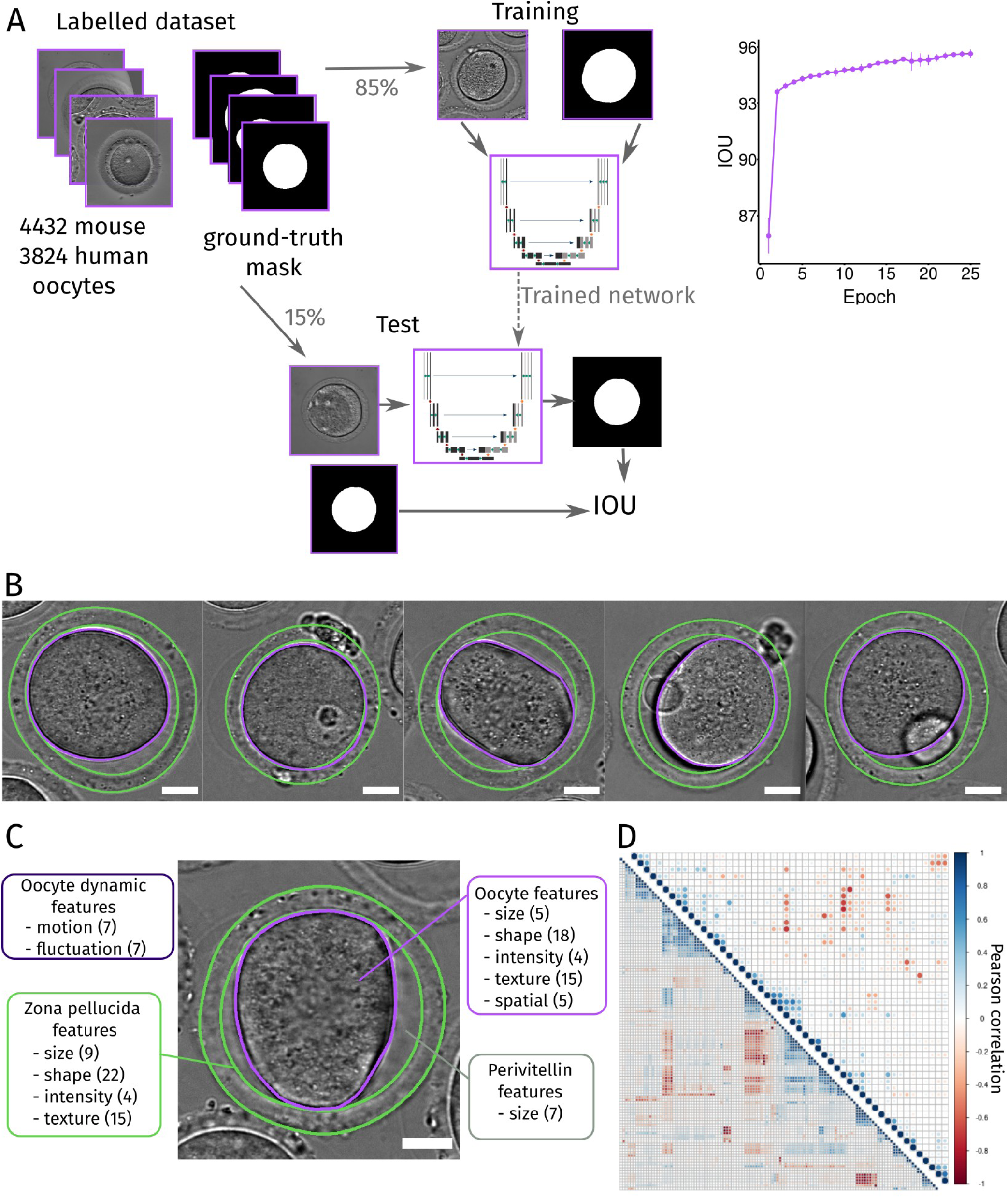
Segmentation and extraction of features from oocytes. A – Scheme of the process used to segment the oocyte contour with neural network. Thousands of mouse and human oocyte images acquired under different conditions (top left panel) with their associated ground-truth (true segmentation mask, top left panel) were split into a training dataset (85% of the images, top middle panel) and a test dataset (bottom panel). Network score is evaluated by the intersection over union (IOU) score during the training iterations (epochs) of the network (top right graph). Once trained, the performance of the network is evaluated by measuring the IOU between its output and the ground truth on the test dataset (bottom panel). B – Examples of segmentation of the oocyte membrane (purple line) and of the zona pellucida contours (green lines) obtained with Oocytor on mouse oocytes at different stages of maturation. Scale bar is 20 µm. C – Features characterizing an oocyte. The number of features appears in parenthesis and the features are grouped by categories: describing the oocyte (purple), its zona pellucida (green), the perivitellin space in between (grey) and the dynamics of the oocyte (dark purple). Scale bar is 20 µm. D – Features correlation and subset selection. Pearson correlation coefficient calculated for each pair of the 118 features (bottom graph) from the values obtained from the training on mouse oocytes dataset. After subset selection (absolute pearson coefficient under 0.75), 49 uncorrelated features were kept (top graph) and used in machine learning algorithms.

### A numerical description of oocytes

To numerically characterize oocytes at a given time point, we defined 118 quantitative measures of their properties (features). These features described the oocyte, its surrounding zona pellucida, its perivitelline space (space between the oocyte and its zona pellucida) and its dynamics (Figure 2C). For this, we used radiomics features (Fornacon-Wood et al., 2020; Rizzo et al., 2018), morphological characteristics that have been shown to be important to discriminate oocyte quality (Inoue et al., 2007; Ozturk, 2020; Rienzi et al., 2011), and other characteristics that could be applied to oocyte description. We implemented the measures of all features based on the segmentation of oocyte and zona pellucida contours from images acquired in transmitted light using our plugin (Supplemental file 1). We defined a high number of features in order to have an unbiased exploration of the oocyte. It is important to note that some features can measure related biological properties and thus have correlated values (Figure 2D, bottom graph). Several methods are available to extract a subset of independent features necessary for some machine learning algorithms (Hall, 2000; Hira and Gillies, 2015). Here, we implemented an unsupervised subset selection based on correlation (Material and Methods, Figure 2D, top graph). Our 118 features or its subset can then be used in machine learning algorithms to classify and characterize oocyte populations.

### A powerful pipeline to automatically phenotype oocytes

In fundamental research, genetic manipulation is often performed to study a particular aspect of oocyte maturation. Our machine learning pipeline allowed to build an algorithm that, after training, should automatically recognize the population of origin of an oocyte. The success/failure of this classification allows to test if the population presents morphological differences and could sort oocytes according to the extent of their mutant phenotype. To demonstrate the potential of our pipeline, we chose to test it on a well-characterized population of Formin-2 knockout oocytes (Fmn2^-/-^) against control mouse oocytes (WT and Fmn2^+/-^) for which we already have datasets of oocytes arrested in prophase I (Al Jord et al., 2021; Almonacid et al., 2015). We kept the smaller dataset to test the performance of our approach, and trained the algorithm with the 2 other combined datasets (Figure 3A). We first verified visually that our segmentation with Oocytor performed well on these new data despite the differences in oocytes phenotypes and modes of image acquisition (Figure S3A). We then applied our pipeline to classify oocytes by their population of origin. For this, in the Analysis step of the pipeline, we compared the performance of different machine learning algorithms (Material and Methods, Figure S3B) that received as input the features describing one oocyte and classified it either as Ctrl or Fmn2^-/-^ (Figure 3A). We obtained the best performance with the Random Forest algorithm (Breiman, 2001) algorithm which was able to recognize control and Fmn2^-/-^ oocytes with approximately 94% fidelity on the training data (Figure 3A, Figure S3B). We tested the performance of this algorithm on an independent set of data and obtained a classification accuracy of about 92% (Figure 3B). Hence, even with a small number of training oocytes (49 control and 49 Fmn2^-/-^ oocytes), our algorithm was able to discriminate oocytes based on their phenotype, thanks to their morphological description.

**Figure 3:**
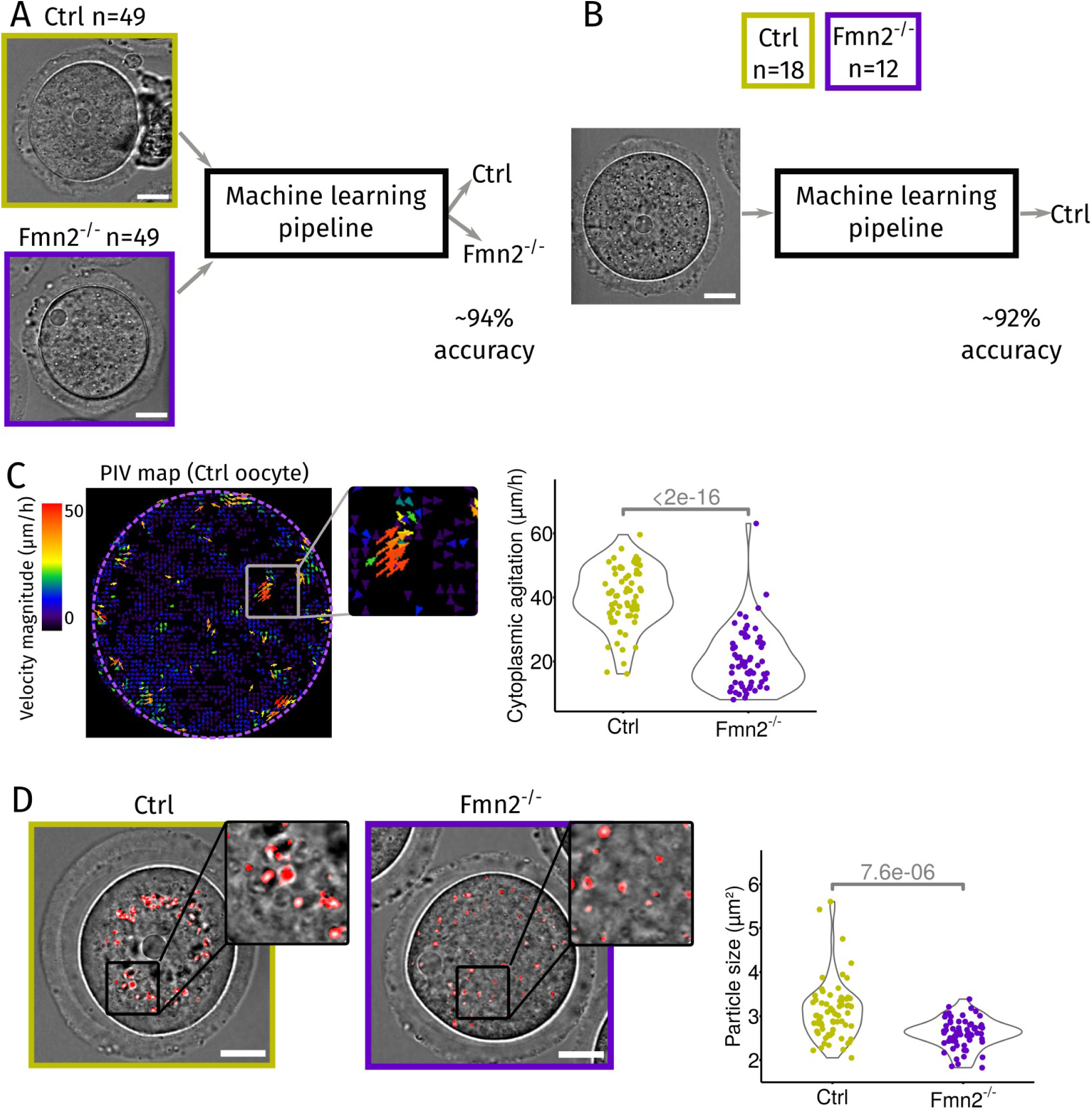
Oocyte phenotyping using our machine learning pipeline. A – Training of our machine learning pipeline to recognize control (Ctrl: WT or Fmn2^+/-^) and Formin-2 knockout (Fmn2^-/-^) oocytes. A training database of 49 Ctrl (dark yellow) and 49 Fmn2^-/-^ (dark purple) oocytes was built from 2 independent datasets and used to train our machine learning pipeline to discriminate Ctrl and Fmn2^-/-^ oocytes. The accuracy of the classification of oocytes as Ctrl or Fmn2^-/-^ was evaluated around 94% (cross-validation on the training data). Scale bar is 20 µm. B – Test of our machine learning pipeline with the selected Random Forest algorithm. A new dataset of 18 Ctrl (dark yellow) and 12 Fmn2^-/-^ (dark purple) oocytes was run through the pipeline and the accuracy of the classification of oocytes as Ctrl or Fmn2^-/-^ was evaluated around 92%. Scale bar is 20 µm. C – Cytoplasmic agitation appears as the most discriminant feature between Ctrl and Fmn2^-/-^ oocytes. Example of a PIV (Particle Image Velocimetry) map extracted from the cytoplasm of a Ctrl oocyte (left panel): arrows indicate the direction of particle motion while their color reflects the magnitude of motion (purple, low to red, high). Insets correspond to a zoom of the image for better visualization. Comparison of the mean value (average from all oocytes and from 5 min duration movies) of the cytoplasmic agitation, measured by the PIV in Ctrl (dark yellow) and Fmn2^-/-^ (dark purple) oocytes (graph on the right panel). Statistical comparison was assessed using a Kolmogorov-Smirnov test (p-value indicated on the graph). D – Cytoplasmic particle size is the second most discriminant feature between Ctrl and Fmn2^-/-^ oocytes. Example of particles detected and measured in our plugin (red dots) in Ctrl (dark yellow box) and Fmn2^-/-^ (dark purple box) oocytes (images on the left panels, scale bar is 20 µm). Insets correspond to a zoom of the image for better visualization. Comparison of the mean cytoplasmic particle size from Ctrl and Fmn2^-/-^oocytes (graph on the right panel). Statistical comparison was assessed using a Kolmogorov-Smirnov test (p-value indicated on the graph).

To understand how the algorithm recognized the two oocyte populations, we examined the weight of the different features (based on the Gini index, Material and Methods). The average cytoplasmic agitation, measured by Particle Image Velocimetry (Tseng et al., 2012), was the most discriminant feature for the algorithm (Figure S3C, Figure 3C). Indeed, cytoplasmic agitation was strongly reduced in Fmn2^-/-^ oocytes (Figure 3C, right panel), confirming previous works showing that cytoplasmic agitation is controlled by the movement of actin vesicles nucleated by Formin-2 (Almonacid et al., 2015; Holubcová et al., 2013). In addition, the second and third most discriminant features were the size and spatial distribution of the cytoplasmic particles (Figure S3C). The difference in spatial repartition most likely reflects the difference in the position of large objects such as the nucleus (centered in control oocytes, off-centered in Fmn2^-/-^ oocytes, Figure 3A, (Almonacid et al. 2015)). The difference in the average particle size (bigger in control oocytes, Figure 3D), a novel feature, could be explained by the difference in cytoplasmic agitation, which tends to center large objects in the cytoplasm (Colin et al., 2020) and thus could favor aggregation of particle into clusters. These results validate and highlight the power of our approach to automatically identify the main differences between two oocyte populations and discover novel characteristics (i.e. particle size) associated to a mutant background.

### Our pipeline predicts mouse oocyte maturation outcome before entry into meiosis

Another major use of our pipeline is its predictive capacity. We used it to predict meiosis entry and the maturation potential of an oocyte arrested in prophase I, before its division. Out of the 468 movies from our dataset of mouse oocyte maturation, 44 oocytes never resume meiosis. We first tested whether we could predict this developmental failure before it happens (Figure 4A). For this, we trained our machine learning pipeline to classify which oocytes would break their nuclear envelope or not, based on the average value of features over the first 12 min of the movies. By cross-validation, we selected the Random Forest algorithm that performed the best for this task (Material and Methods, Figure S4A). Finally, we trained our selected algorithm on all the data and tested its performance on a new independent dataset of 69 oocytes, 7 of which did not resume meiosis. We obtained a balanced accuracy of about 91% (Figure S4A, right). Therefore, our algorithm has the power to correctly predict which prophase I oocytes will fail to break their nuclear envelope. The most discriminant features associated with meiosis entry were the thickness and texture of the zona pellucida and the size of the oocyte-zona pellucida complex (Figure S4C). Oocytes that remained arrested in prophase I were smaller, with a thin and heterogeneous zona pellucida (Figure 4C, red), consistent with them not finishing their growth (Wassarman and Litscher, 2013), further validating our approach.

**Figure 4:**
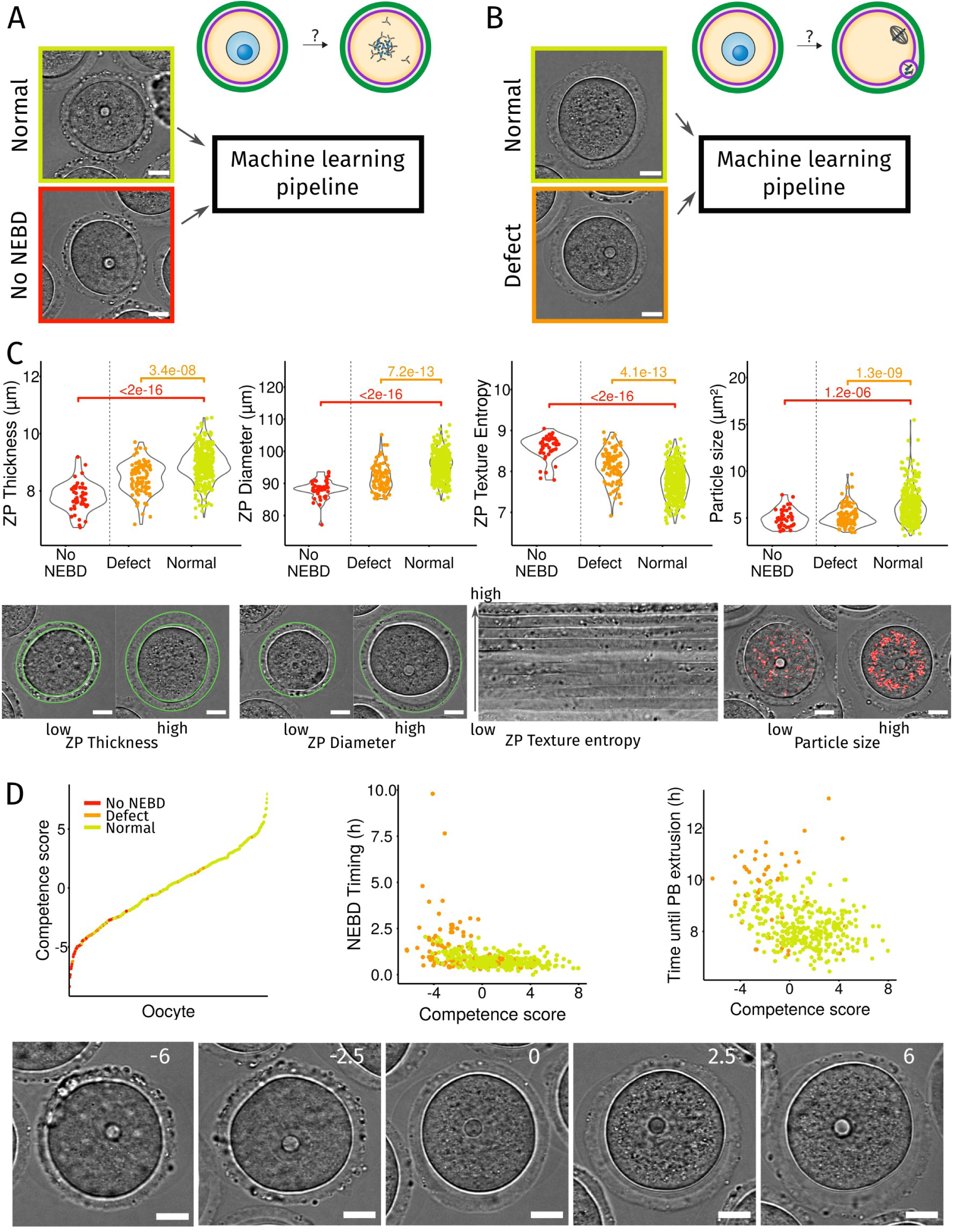
Prediction and characterization of mouse oocyte maturation with our machine learning pipeline. A – Prediction of failure to enter meiosis I. Our machine learning pipeline is trained with our mouse oocyte maturation dataset. Each oocyte was labelled as Normal if it succeeded to break its nuclear envelope (yellow box, 424 oocytes) and No NEBD otherwise (red box, 44 oocytes). B – Prediction of a defect in maturation after meiosis resumption. Each oocyte was labelled as Normal if no defect was detected (yellow box, 327 oocytes) and as Defect if the oocyte did not extrude a first polar body, resorbed its polar body, entered meiosis I after an abnormal delay or extruded its polar body after a delay (orange box, 96 oocytes). C – Discriminant features for NEBD failure or maturation defect. Comparison of the values of the discriminant features for oocytes that do not enter meiosis I (red, No NEBD), with maturation defect (orange, Defect) or normal (yellow, Normal). P-value indicated on the graphs were calculated with a Kolmogorov-Smirnov test. From left to right panels: graphs show the comparison of the average thickness of the zona pellucida (average distance between the inner and the outer contours), average value of the zona pellucida outer diameter, average value of the entropy of the texture of the zona pellucida (measured by Haralick’s entropy), and average cytoplasmic particle size. Illustration of oocytes with low and high values of the discriminant features (bottom panel). The ZP contours are highlighted in green, cytoplasmic particles in red. Scale bar is 20 µm. D – Quantitative measure of oocyte competency. The competence score is defined as a linear combination of the 4 features presented in C: competence score = ZP outer perimeter + ZP texture entropy + ZP thickness + Cytoplasmic particle size. Sorted competence scores from all oocytes of the mouse maturation dataset (graph on the top left panel). Each point represents one oocyte, and the color the category to which the oocyte belongs: no NEBD (red), maturation defect (orange), normal (yellow). Timing of NEBD according to the oocyte competence score (graph on the top middle panel). Time spent between NEBD and first polar body extrusion according to the oocyte competence score (graph on the top right panel). Examples of oocytes observed in transmitted light presenting different competence scores indicated in white on the images (bottom panel), increasing from left (very low competence score) to right (high competence score). Scale bar is 20 µm.

Next, we aimed to predict and characterize the potential of oocytes to properly mature (Figure 4B). We analyzed only the movies of oocytes that underwent nuclear envelope breakdown and temporally aligned them based on the NEBD timing calculated with Oocytor. We considered that oocytes had maturation defects when they did not extrude or when they resorbed their first polar body (PB), or if the timing of NEBD or first polar body extrusion was abnormally late (Figure S5). We trained machine learning classifiers to predict oocytes that will have a maturation defect based on the average values (over 12 min) of the features taken 15 min before NEBD. Our selected algorithm (balanced Random Forest, Figure S4B) had a balanced accuracy of nearly 80% on the training data (Figure S4B) and 90% on the test dataset (which contained 3 out of 62 oocytes with maturation defects). Thus, our approach allowed to predict the maturation potential of oocytes before maturation even resumes.

### Oocyte quality can be scored from 4 non-dynamic features

The two most discriminant features used by the algorithm for prediction of oocyte maturation outcome were the zona pellucida texture and the cytoplasmic particle size (Figure S4C). Zona pellucida texture (measured by Haralick’s entropy, see Supplemental File 1) was a determinant feature of oocyte quality for both the entry and the success of oocyte maturation, making it highly relevant for assessing oocyte potential. This feature reflects the heterogeneity of the zona (Figure 4C, third panel). It was highest in oocytes that did not start maturation (Figure 4C, third panel, red) and was relatively high in oocytes that had maturation defects (Figure 4C, third panel, orange). The second discriminant feature for successful oocyte maturation was cytoplasmic particle size, which was smaller in oocytes that failed to resume or to properly end meiotic maturation (Figure 4C, fourth panel, red and orange). At last, both the thickness of the zona pellucida and the size of the oocyte-zona pellucida complex were good indicators of oocyte quality (Figure 4C, first and second panels). Thus, the characteristics of the zona pellucida are a good measure of oocyte maturation potential.

To evaluate the quality of an oocyte, we defined a score based on these most relevant features for resuming and completing maturation. This competence score, an indicator of oocyte quality, which corresponds to its capacity to develop properly, was defined as a linear combination of the standardized values of the 4 most discriminant features: texture and thickness of the zona pellucida, size of cytoplasmic particles and perimeter of the outer limit of the zona pellucida (Figure 4D and Material and Methods). Importantly, this score was based on non-dynamic features and therefore only required a single image acquisition of an oocyte arrested in prophase I. Consistently, most oocytes that did not resume meiosis had a low competence score (Figure 4D, left panel, red), whereas oocytes that matured properly had an overall high competence score (Figure 4D, left panel, yellow). Moreover, the competence score correlated with the maturation dynamics of oocytes that entered meiosis I: oocytes with low score resumed meiosis more slowly (Figure 4D, middle panel, Pearson coefficient r=-0.4, p-value<10^-16) and were delayed in first PBE extrusion (Figure 4D, right panel, Pearson coefficient r=- 0.36, p-value=4.8 10^-13). Hence, our competence score allows to rank oocytes arrested in prophase I by their developmental potential (Figure 4D bottom panel).

### Heterogeneity of the zona pellucida is predictive of *in vitro* maturation of human oocytes

As our algorithm proved to accurately estimate the quality of mouse oocytes, we then tested whether the same discriminating features would be applicable to human oocytes. For this, we used a dataset of 72 human oocyte maturation movies acquired in clinics (Materials and Methods). Oocytes were collected from patients after hormonal stimulation, and those that did not complete maturation, and therefore inappropriate for intracytoplasmic sperm injection (ICSI), were followed overnight for research purposes. The dataset was a mix of prophase and meiosis I oocytes at the beginning of the movies, hence it could not be temporally aligned as for mouse oocytes, increasing the variability. 48 out of 72 of these oocytes eventually succeeded in extruding a first polar body. To test whether our previously identified features could predict polar body extrusion, we used our plugin on these movies. Since we trained our neural networks for cortex and zona pellucida segmentation on both mouse and human oocytes, we achieved a good performance on the segmentation of human oocytes despite the presence of obstacles in the field of view, such as for example the presence of cumulus cells (Figure S6A). This demonstrates that Oocytor can successfully segment human oocyte and zona pellucida contours. With Oocytor, the values of our 118 features were then extracted. To visualize the distribution of the values of these features at the beginning of the movies, the U-MAP reduction technique (McInnes et al., 2018) was applied. No strong morphological differences were detected between extruding and non-extruding oocytes (Figure S6B left panel). We then compared extruding and non-extruding human oocytes using the values of the discriminant features identified from mouse oocytes at the beginning of the movies (Figure S4C). The differences were not significant, except for the texture of the zona pellucida (Figure S6B middle and right panels). As those movies could not be temporally aligned, the analysis was performed on the values of the features at the beginning of the movies, hence oocytes were at different stages of development contrary to our analysis on mouse oocytes, which increases the variability of the features values. Moreover, it is important to remind here that the population of human oocytes analyzed in these experiments correspond to oocytes not responding to the hormonal stimuli and thus probably of lower quality. Hence this population may not be comparable to the mouse one and may not represent a physiologically relevant cohort of human maturing oocytes. Consistent with this, the nucleus visible at the beginning of 39 movies, was generally not centered, a feature known to correlate with defects in polar body extrusion (Levi et al., 2013).

Nevertheless, this analysis of low quality human oocytes before metaphase II, the only ones accessible sor far for research purpose, showed the importance of the texture of the zona pellucida: a more heterogeneous zona correlated with maturation defects both in human and mouse oocytes. This feature has also been reported as an important marker of quality of human matured oocytes and human embryos, to assess their potential to develop up to the blastocyst stage and their implantation success (Rienzi et al., 2011). In particular, it was shown that the optical birefringence measured by polarized light microscopy was higher and more uniform in human oocytes that developed properly (Montag and van der Ven, 2008). Altogether, our results suggest a critical role of the zona pellucida in oocyte maturation and revealed that its morphology can reflect the quality of the oocyte even before maturation.

## Discussion

We developed a computational tool to segment and describe numerically oocytes, associated with a machine learning pipeline to phenotype oocyte populations or predict and characterize oocyte development. Our tool was setup on images acquired in transmitted light which allows its use both in academia and in clinics. As all the steps of the pipeline are automatized, our tool can be used to screen an important number of oocytes for large scale studies.

To make our tool user friendly, we implemented it as a Fiji plugin (Schindelin et al. 2012), Oocytor, available as an open-source on Github (Material and Methods). Oocytor can perform 3 tasks on label-free images: segmentation of the oocyte and its zona pellucida contours, NEBD detection and extraction of hundreds of morphological features. For the two first tasks, we took advantage of the power of deep learning algorithms and trained U-Net like networks. One of the pitfalls of machine learning algorithms and neural networks in particular, is their major dependence on the data used to train them. In general, these networks perform very well on a dataset close to the training one, but can fail completely on a new one. We attempted to produce more robust and versatile networks by feeding them with a variety of images (Möckl et al., 2020), pooling human and mouse oocytes images from different projects, acquired in different laboratories and clinics. Nonetheless, the networks could still yield limited results on oocytes never seen before by the networks, such as, for example, oocytes from a different species. In that case, a new network could be trained with only few images from these “foreign” oocytes, by using the transfer learning technique (Bengio, 2012) on our neural network available in the Github repository. Oocytor also offers the possibility to automatically define the NEBD timing of mouse oocytes, thus avoiding manual annotation and providing a temporal reference to align maturation movies. This property can turn-out to be a true time-saver for studies implemented on large cohorts of oocytes. We implemented the measure of 118 features, allowing to numerically describe an oocyte at any stage of its development. This list could be further expended. In particular, it would be interesting to add segmentation and features to better describe polar body properties, which certainly would help a better characterization of metaphase II arrested oocytes. New neural networks could also be trained to infer the position of the nucleus, spindle, or other organelles from images acquired in transmitted light using data coupled with fluorescence labeling (Christiansen et al., 2018; Ounkomol et al., 2018; Wieslander et al., 2021). This approach is very promising for fundamental research to avoid fluorescence labeling. These steps of synchronization, segmentation and numerical description are independent and can then be used to answer a variety of fundamental research studies on oocytes.

Based on the numerical description of oocytes, we developed an interpretable machine learning pipeline to classify oocytes and identify major differences between oocyte populations. Here, we demonstrated that our algorithm can discriminate oocytes from different genetic backgrounds, such as control and Fmn2^-/-^ oocytes, after training with only a small dataset, and revealed main morphological differences between these two populations. This phenotyping could be very useful in the mouse oocyte community where genetic modifications are often used as tool to identify gene function without knowing all the consequences on oocyte characteristics. We thus propose here a new approach to systematically identify new differences in an unbiased and automatic manner and thus to potentially reveal novel gene function during oocyte development.

We then used this approach to explore oocyte quality within a wild-type population for clinical applications. We developed our approach on mouse oocytes, allowing to have access to more data and possible manipulations. We successfully trained an algorithm to recognize oocytes that failed to resume meiosis. These oocytes were smaller with a thin and heterogeneous zona pellucida. It has been previously shown that small size is a sign of oocyte incompetence, which corresponds to oocytes unable to successfully develop up to the metaphase II stage (Hirao et al., 1993; Kanatsu-Shinohara et al., 2000; Wickramasinghe et al., 1991). This small size is most probably an indicator of unfinished oocyte growth. The zona pellucida thickness correlated with the size of the oocyte before NEBD in our mouse data (Pearson correlation r=0.42, p-value<10^-16), consistent with the fact that the zona pellucida thickness and the oocyte diameter increase concomitantly during oocyte growth (Wassarman and Litscher, 2013). However, this correlation was lost after NEBD (Pearson correlation at NEBD+5h r=0.18, p-value=1.6 10^-3), the zona pellucida thickness remaining constant while the oocyte size decreased. Therefore, ZP thickness brings an information on the maturation potential of the oocyte which is different from the oocyte size. In human oocytes, no correlation has been found between zona pellucida thickness and oocyte diameter in metaphase II-arrested oocytes (Bertrand et al., 1995), consistent with our findings of loose correlation in mouse oocytes. Moreover, in human embryos, zona pellucida thickness was positively associated with development up to the blastocyst stage and to embryo quality (Høst et al., 2002; Raju et al., 2007; Shen et al., 2005), but was negatively correlated with fertilization success (Bertrand et al., 1995, Marco-Jiménez et al., 2012). Thus, zona pellucida thickness could be a relevant feature for assessing oocyte (our study) and embryo quality.

We trained an algorithm to predict maturation defects of mouse oocytes before NEBD. Our machine learning algorithm revealed that mouse oocytes with maturation defects had a more heterogeneous zona pellucida and smaller cytoplasmic particles (Figure 4C). The heterogeneity of the zona pellucida, measured by Haralick’s entropy, was a discriminant feature for both the initiation and the success of maturation. Moreover, this feature was also relevant for predicting human oocyte maturation potential. This texture could be related to the birefringence of the zona pellucida, which is used to assess human embryo quality (Gabrielsen et al., 2000; Ozturk, 2020; Raju et al., 2007; Rienzi et al., 2011) and the reflect the structuration of the zona. Cytoplasmic particles were smaller in oocytes with a low competence but also in Fmn2^-/-^ oocytes (Figure 3D), so this difference observed in the control oocyte population could be related to differences in the cytoplasmic actin activity that was reduced both in oocytes of low competence score and in Fmn2^-/-^ oocytes (Figure S5B, Figure 3C). Previous studies showed that actin vesicle activity follows a gradient from the cortex to the center, generating a non-specific centering force (Almonacid et al., 2015; Colin et al., 2020), that could favor cytoplasmic particle aggregation. This reduced activity, correlating with a low maturation potential (Figure S5B), was also associated with slower maturation dynamics (Figure 4D, middle and right panels).

To conclude, our machine learning pipeline could predict oocyte maturation with a fidelity between 80% and 90%. Furthermore, we identified two new features, the texture of the zona pellucida and the size of cytoplasmic particles, as good predictors of mouse oocyte maturation potential. Moreover, we showed that the zona pellucida texture could be a good predictor of human oocyte maturation potential. We also defined a competence score to rank mouse oocytes by their capacity for maturation and revealed a defect in the cytoplasmic activity in poor quality oocytes. This ranking could be used to improve *in vitro* maturation protocols by identifying the most promising oocytes. By adding new features to Oocytor, such as polar body contour detection, it might also be possible to adapt our approach to assess the potential of oocytes for fertilization and embryonic development.

## Material and methods

### Dataset

#### Mouse oocyte maturation

Our study was primarily based on a dataset of 468 movies of *in vitro* matured mouse oocytes. An additional dataset of 69 mouse oocyte maturation movies were acquired independently, after the pipeline development, and used only to test its final performance.

OF1 or C57Bl6j female mice (Charles River laboratories) aged between three and ten weeks were used for experiments. The prophase I-arrested oocytes were collected according to laboratory protocol in M2 medium + Bovine Serum Albumine (BSA, A3311, Merck) supplemented with 1µM Milrinone (Reis et al., 2006) (M4659, Merck). For *in vitro* maturation, the oocytes were washed out from Milrinone and cultured in M2 medium under mineral oil (N8410, Merck) at 37°C. They were then placed under a Leica DMI6000B microscope equipped with a Plan-APO 40x/1.25 NA oil immersion objective, a motorized scanning deck and an incubation chamber (37°C), a Retiga 3 CCD camera (QImaging, Burnaby) coupled to a Sutter filter wheel (Roper Scientific), and a Yokogawa CSU-X1-M1 spinning disk. Images were acquired using Metamorph (Universal Imaging, version 7.7.9.0) every 3 min in transmitted light with the objective 20x/0.75 NA for twenty hours at 37°C.

#### Fmn2^-/-^ dataset

We re-used datasets of control and Fmn2^-/-^ genotypes of oocytes arrested in prophase I from previous studies (n=25 Fmn2^+/-^, n=12 Fmn2^-/-^ (Al Jord et al., 2021); n=18 WT, n=12 Fmn2^-/-^ (Almonacid et al., 2015)) and we performed a supplemental experiment with oocytes from both genotypes (n=24 WT and n=37 Fmn2^-/-^) to increase the training dataset (see above protocol). Oocytes were imaged every 1 min (movies were under-sampled for dataset when imaging frequency was higher) for 5 min or more. Datasets were not acquired with the same microscope objective resolution (0.1135 µm/pixel, 0.1613 µm/pixel and 0.227 µm/pixel) so we resized the images to a resolution of 0.1613 µm/pixel to allow direct comparison.

#### Human oocyte maturation

Human oocyte maturation movies were acquired at the Cochin hospital for research purposes (72 movies). Immature human prophase I-stage and meiosis I-stage oocytes found in cohorts retrieved for the purpose of intracytoplasmic sperm injection (ICSI) can be used for research according to the French legislation, with the consent from patients. The *in vitro* maturation study was performed with oocytes (n=72) that were donated for research by patients undergoing ART protocols. It was approved by the Germetheque Biobank (BB-0033–00081) under the number 20160912. Once the ICSI procedure was performed for the patients, oocytes that did not reach metaphase II were included into the protocol. Culture dishes were prepared, covered with mineral oil (Irvine Scientific, Ireland), warmed and pre-gassed before *in vitro* maturation. Cumulus cells had been removed from the oocytes by brief treatment with hyaluronidase IV-S (Sigma-Aldrich, USA) at 37°C. These desynchrionized oocytes were incubated in Continuous Single Culture Complete medium (CSCM-C; Irvine Scientific) in an embryoscope (Geri time-lapse system, Genea Biomedx Pty Ltd) at 37°C, 6% CO2, 5%O2 to record the maturation process. *In vitro* maturation to the NEBD, metaphase I, metaphase II stages, activation or atresia was evaluated 24 hours later. Oocytes were considered as ‘matured’ when the first polar body was present.

#### Additional datasets

##### Mouse datasets

To train our neural network for segmentation, we also used data from other projects to diversify our training. We used movies of mouse oocyte maturation acquired in transmitted light for wild-type oocytes as well as extra-soft oocytes from a previous study (Bennabi et al., 2020). To train our neural network to determine the timing of NEBD, we also used another dataset of mouse maturing oocytes where the NEBD timing had already been manually annotated (120 movies).

##### Human datasets

Through the ICSI procedure at the Cochin hospital (see above), we also had access to images of metaphase II oocytes that we used to diversify our training data (658 images). Moreover, we also added images from movies following the development of fertilized oocytes from the Hôpital Femme Mère Enfant in Lyon (551 movies). The cumulo-oocyte complexes (COCs) were obtained after transvaginal follicular puncture of patients treated for infertility. One hour after COCs retrieval, they were stripped by enzymatic digestion (hyaluronidase®; CooperSurgical, Malov, Denmark) and by a mechanical action. The stripped oocytes were fertilized by intracytoplasmic sperm injection (ICSI). After ICSI, the fertilized oocytes were immediately cultured in oil-coated Cleav® medium (CooperSurgical) in a time-lapse system (EmbryoScope®; Vitrolife, Viby, Denmark). The culture was monitored by successive image acquisitions (1 image every 15 minutes).

### Ground-truth database creation

Before training the neural networks, we tried to segment images using an approach based on thresholding and morphology. However, the results were highly dependent on the input parameters and could not be used without manual validation and correction. We therefore chose to use a deep learning approach instead to improve the performance. This preliminary segmentation allowed to build an initial database with the manually validated and corrected ground truth segmentation. Moreover, in the additional dataset coming from a previous project (Bennabi et al., 2020), some oocytes were also stained with a membrane fluorescent marker, which allowed to have a direct access to the ground-truth images for these oocytes. In the end, we created a database of 8256 images (4432 mouse, 3824 human) with their associated ground-truth.

Eventually, the creation of the ground-truth for the detection of the zona pellucida was more challenging since the contrast was often very low. To achieve this, the segmentation was done or corrected manually on a large part of the dataset, by defining an ellipse around the zona pellucida, and thus lacked precision. This could explain the lower performance of the neural network on zona pellucida segmentation against this imprecise ground-truth. The database for the zona pellucida consisted of 3578 images (2361 mouse, 1217 human) with their associated ground-truth.

It is important to note that multiple images per movie were used in the datasets to increase the number of images. However, when a dataset was split between training and test subsets, as well as when the training dataset was split for cross-validation, images extracted from the same movie were always in the same subset, to ensure that we had independent datasets and avoid data leakage (Wen et al., 2020).

### Machine learning pipeline

#### Pipeline preprocessing

First, movies/images had to be preprocessed. When several oocytes were present in the same movie, they were automatically cropped into several image stacks of a single entire oocyte. Moreover, the movies were aligned to keep the oocyte at the same position in the images for measures of dynamic features. Finally, intensity normalization was performed to homogenize the images coming from different sources.

#### Oocytor plugin

Oocytor is a Fiji plugin (Schindelin et al. 2012), implemented in Java. The source code can be found on Github (https://github.com/gletort/Oocytor), as well as a compiled version presented as a ready to use plugin, with installation instructions. Oocytor can perform 3 tasks: oocyte contour segmentation, NEBD detection and features extraction. Segmentation and NEBD detection are based on our neural network trained on large databases in python. To run the already trained neural network in Fiji, Oocytor uses CSBDeep plugin (http://sites.imagej.net/CSBDeep) (Weigert et al., 2018). To calculate some features, Oocytor uses FeatureJ (http://imagescience.org/meijering/software/featurej). Several macros are also proposed in the Github repository to facilitate Oocytor usage with several data folders.

#### Oocytor NEBD detection

##### Neural network implementation and training

we built our neural network to classify images based on the presence of a nucleus or not with the VGG-16 architecture (Simonyan and Zisserman, 2015) and tested variations around this architecture. The final architecture is shown in Figure S1A. To train the neural network for NEBD detection, we used a one-time data augmentation, by flipping the images, on the training images. Images were resized to 256*256 pixels and normalized before being fed to the neural network. We trained the neural networks for 25 epochs, with batch normalization and a batch size of 30, with ReLU activation functions. To adjust the network hyper-parameters, we performed 10-fold cross-validation on the training dataset (Figure S1B, selection of the number of filters n in the initial layer). The selected network was then trained on the full training dataset and its performance was tested on the independent test dataset (Figure 1C right panel).

##### NEBD determination in Oocytor

each image of the movie is resized to 256*256 pixels and normalized before running through the neural network. This gives a score of probability of absence of a nucleus at each time point (Figure 1D). The evolution of this score according to time is locally smoothed and NEBD is calculated as the first transition point from low to high score (Figure D bottom left panel).

### Oocytor segmentation

#### Neural network implementation and training

We based our neural network architecture for cortex and zona pellucida segmentation on the U-Net architecture (Ronneberger et al., 2015). We tested several configurations around this architecture (number of layers, activation function, variation of the architecture (Falk et al., 2019; Gadosey et al., 2020; Ibtehaz and Rahman, 2020)) and opted for a classical U-Net architecture, shown in Figure S2A. Note that we could have trained a single network to segment at the same time the zona pellucida and the cortex, which can slightly improve the performance of the network (Firuzinia et al., 2021). However, we preferred to keep two independent networks for more flexibility. Images were resized to 256*256 pixels and normalized before being fed to the neural networks. We trained the neural networks for 25 epochs, with batch normalization and a batch size of 30, with ReLU activation functions. To adjust the network hyper-parameters, we performed 6-fold cross-validation on the training dataset (Figure S2B-C, selection of the number of filters n in the initial layer). Based on these results, we selected a U-Net like architecture with n=8 initial filters for cortex segmentation (Figure S2B) and with n=16 initial filters for zona pellucida segmentation (Figure S2C). The selected networks were then trained on the full training dataset and their performance were tested on the independent test dataset (Figure S2D).

#### Implementation of segmentation

Oocytor first resizes (to 256*256 pixels images) and normalizes input images or movies. To increase the robustness of the plugin, the results of two neural networks trained on the same task (segmentation of oocyte contour or zona pellucida boundaries) were combined (Figure S2E, “Get cortex” function of Oocytor). The resulting binary images are converted into Fiji ROIs that can eventually be refined to local variation of intensity or smoothed, depending on input parameters of the plugin.

### Feature selection

We calculated the Pearson correlation between all features (Figure 2D) and set a threshold (0.7 here) above which features were considered correlated. By iterations, we kept the features with the most connections (correlated features) and removed those connected to it, until we obtained a subset of uncorrelated features that were below our threshold (Figure 2D, top).

#### Feature standardization

Standardization was performed to shift features values in a similar range for all features. For each feature, the mean and standard deviation (std) were calculated from the training data. The feature values were then updated as: f = (f-mean)/std.

### Machine learning methods

After features extraction with Oocytor, machine learning analysis were performed with the R software (R Development Core Team, 2008). We used the “randomForest” and “e1071” packages (Liaw and Wiener, 2002; Meyer et al., 2019) for the classification algorithms tested. UMAP projections were calculated with the “umap” package (Konopka, 2020). Finally, graphs were generated with the “ggplot2” package (Wickham, 2016).

#### Classification methods

we tried 3 standard machine learning methods in the Analysis step: Naive Bayes classifier (Rish, 2001), Support Vector Machine (Vapnik et al., 1996) and Random Forest (Breiman, 2001). Each of these algorithms received as input the features describing one oocyte and classified it. We used cross-validation techniques to measure the performance of our training and to select the best method (Figure S3B, Figure S4A-B). The selected method was finally trained on all the training data and its performance was always tested on an independent dataset.

#### Data imbalance

Datasets were strongly imbalanced between oocytes that entered maturation or not (Figure S4A left) as well as between oocytes that matured correctly or not (Figure S4B left). As we were interested in building a tool that would discriminate them based on images and not by considering the frequency of each class, we equilibrated the training datasets and measured the balanced accuracy to score the classification (Figure S4A middle and Figure S4B middle). For this, we used the number of the smallest dataset for both classes in the classification algorithm (under-sampling). We also tried to balance the dataset by data augmentation (oversampling) of the smallest dataset with the SMOTE technique (Chawla et al., 2002) but did not obtain better results (not shown).

#### Performance score

the score used to measure the performance of the algorithms were:

- accuracy = (TP+TN)/(TP+TN+FP+FN)
- Recall, True Positive Rate (TPR) = TP/(TP+FN)
- Precision = TP/(TP+FP)
- True Negative Rate (TNR) = TN/(TN+FP)
- Balanced accuracy = (TPR+TNR)/2

where TP, TN, FP and FN correspond respectively to: True Positive, True Negative, False Positive and False Negative.

#### Feature importance

to assess the contribution of each feature in the Random Forest algorithm, we considered the Gini index of the features in the decision trees. A higher Gini index indicates that the feature provides stronger separation of the population in the tree. Thus, features with higher Gini index were the most discriminant in the algorithm.

## Acknowledgments

We thank Maria Almonacid, Adel Al Jord, Isma Bennabi and Flora Crozet for sharing experimental data from their previous work; Lucie Barbier and Jean-Yves Tinevez for comments on the manuscript; all members of the Terret-Verlhac laboratory and of the CIRB imaging facility Orion for discussions.

## Competing interests

No competing interest to declare.

## Supplemental Figures

**Figure S1:**
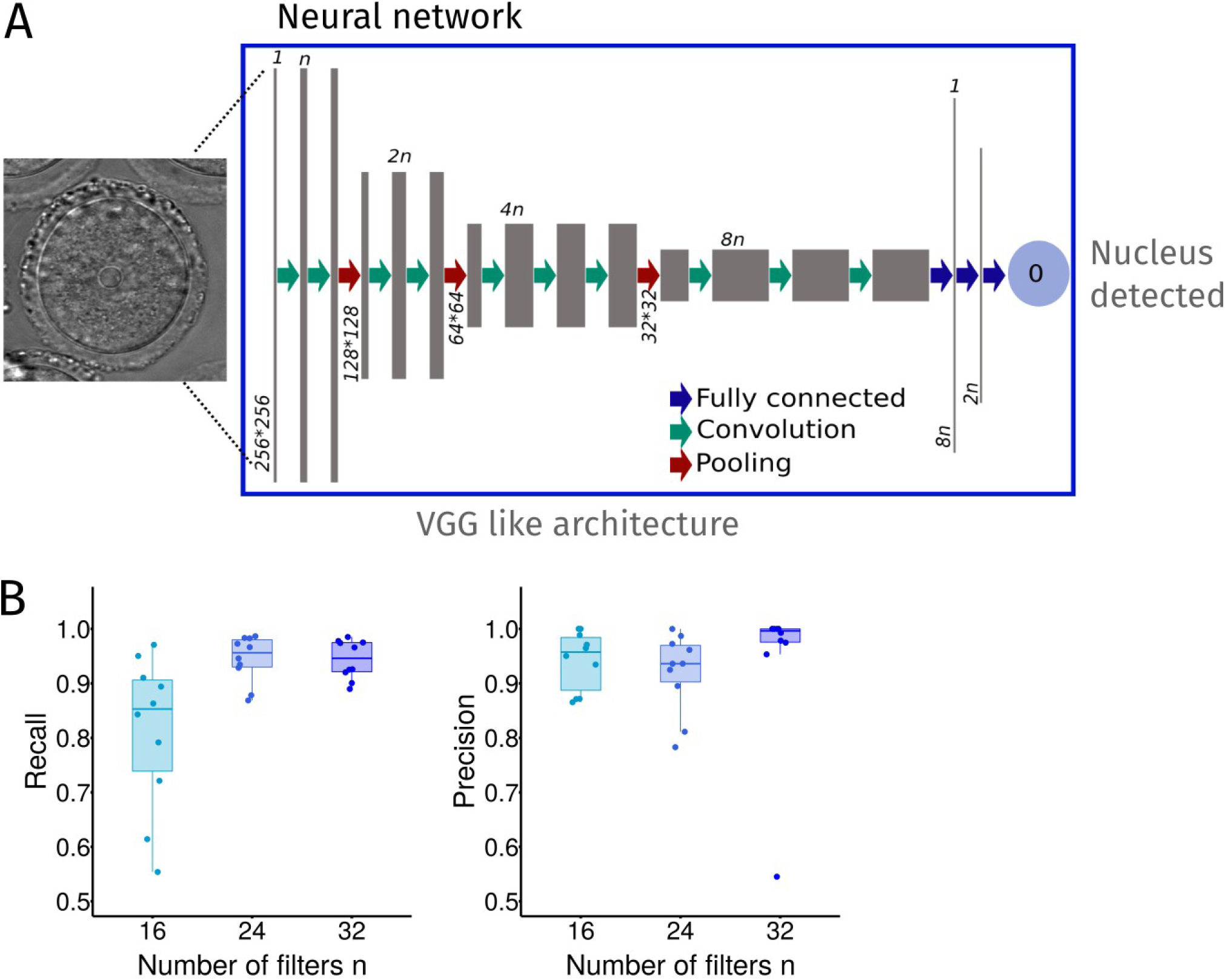
Neural network architecture for nucleus detection. A – Architecture of the selected neural network, based on the VGG-16 architecture. The network takes as input one image of an oocyte and gives as output a score of detection for the nucleus (0: presence of a nucleus, 1: absence). The first convolution layer applies (n) filters, which controls the size of the network and we tested several values for this parameter. B – Score of nucleus detection from neural networks for different values of the number of initial filters (n) controlling the network size. The detection score was evaluated by Recall: TP/(TP+FN) and Precision: TP/(TP+FP), where TP=True Positive, FN=False Negative, FP=False Positive. The selection of the best parameter (n) was done with a 10-fold cross validation technique on the training dataset: data are split in 10 subsets, 9 are used for training, and the score is evaluated on the remaining one. The operation is repeated 10 times so that each subset is used as the test subset once. The final selected network had n=32 initial filters.

**Figure S2:**
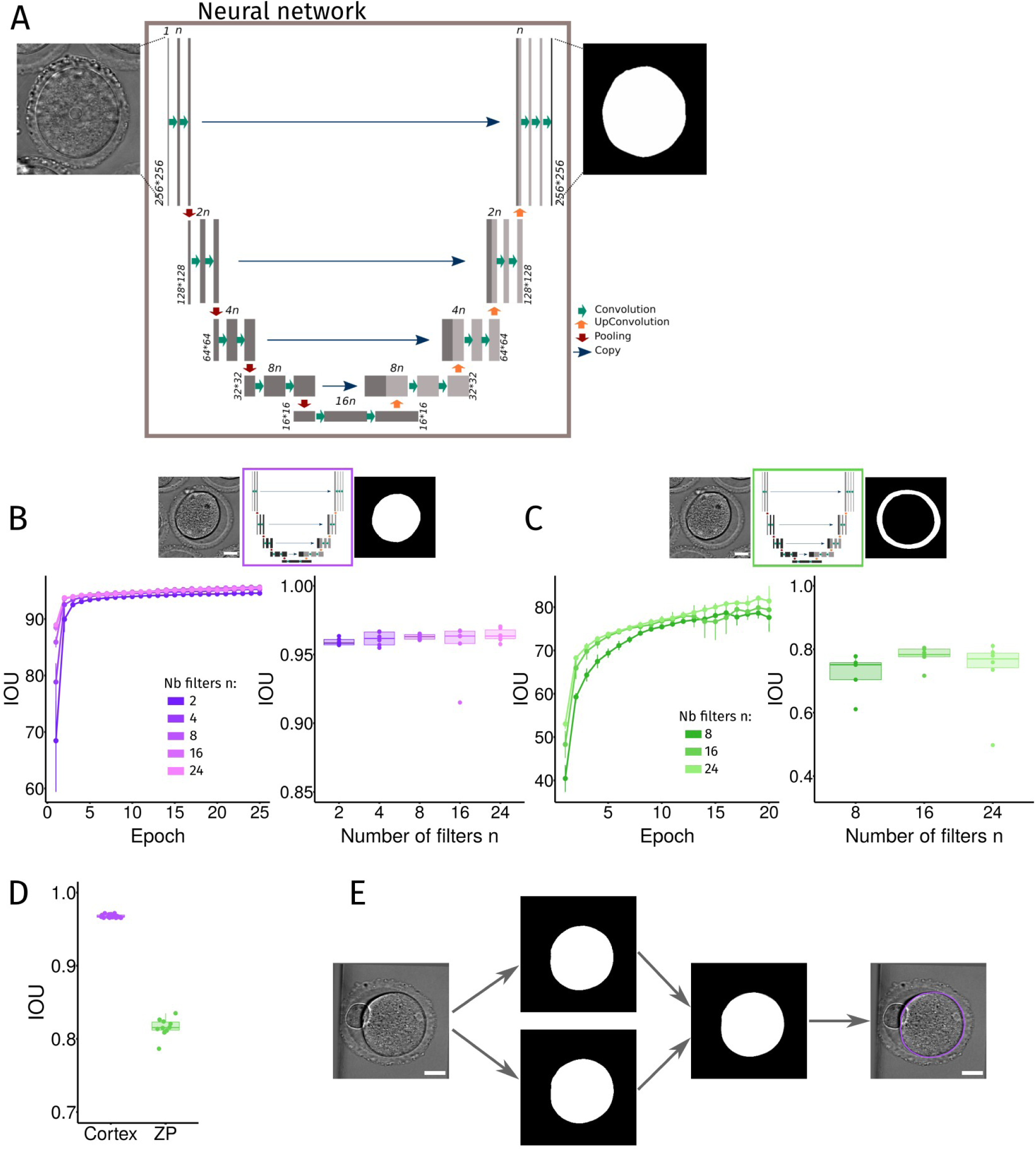
Neural network architecture for the detection of the oocyte contour. A – Architecture of the selected neural network, based on a U-Net architecture. The network takes as input one image resized to 256*256 pixels of an oocyte and outputs a binary mask with the interior of the oocyte. The size of the network is controlled by (n), the number of filters in the first convolution layer. B – Selection of the proper neural network by cross-validation for oocyte membrane segmentation (purple box). The training dataset was split in 6 subsets for cross-validation. Networks with n=2, 4, 8, 16 or 24 initial filters were trained on 5 of the 6 subsets. Their performance during training was evaluated at each iteration (epoch) by the intersection over union (IOU) score between the network outputs and the ground truth images (left graph). The final performance of the trained network was validated on the respective remaining subsets by the IOU score (right graph). The network with n=8 filters was selected. Scale bar is 20 µm. C - Selection of the proper neural network by cross-validation for zona pellucida contours segmentation (green box). By 6-fold cross-validation, the performance of neural networks with n=8, 16 or 24 initial filters (n) was evaluated during training by the IOU score (left graph) and after training on the remaining test subset (right graph). The network with n=16 filters was selected. Scale bar is 20 µm. D – Performance of the neural network for membrane (purple dots) and zona pellucida (green dots) segmentation on independent test dataset for mouse oocytes measured by the IOU score. E – Segmentation of the oocyte membrane with Oocytor. The input image is run through 2 neural networks trained on all our training dataset (mouse and human oocytes). Two networks were used to increase the robustness. Their outputs were combined in a final binary mask used to determine the oocyte contour. Scale bar is 20 µm. The procedure is the same in the case of zona pellucida segmentation.

**Figure S3:**
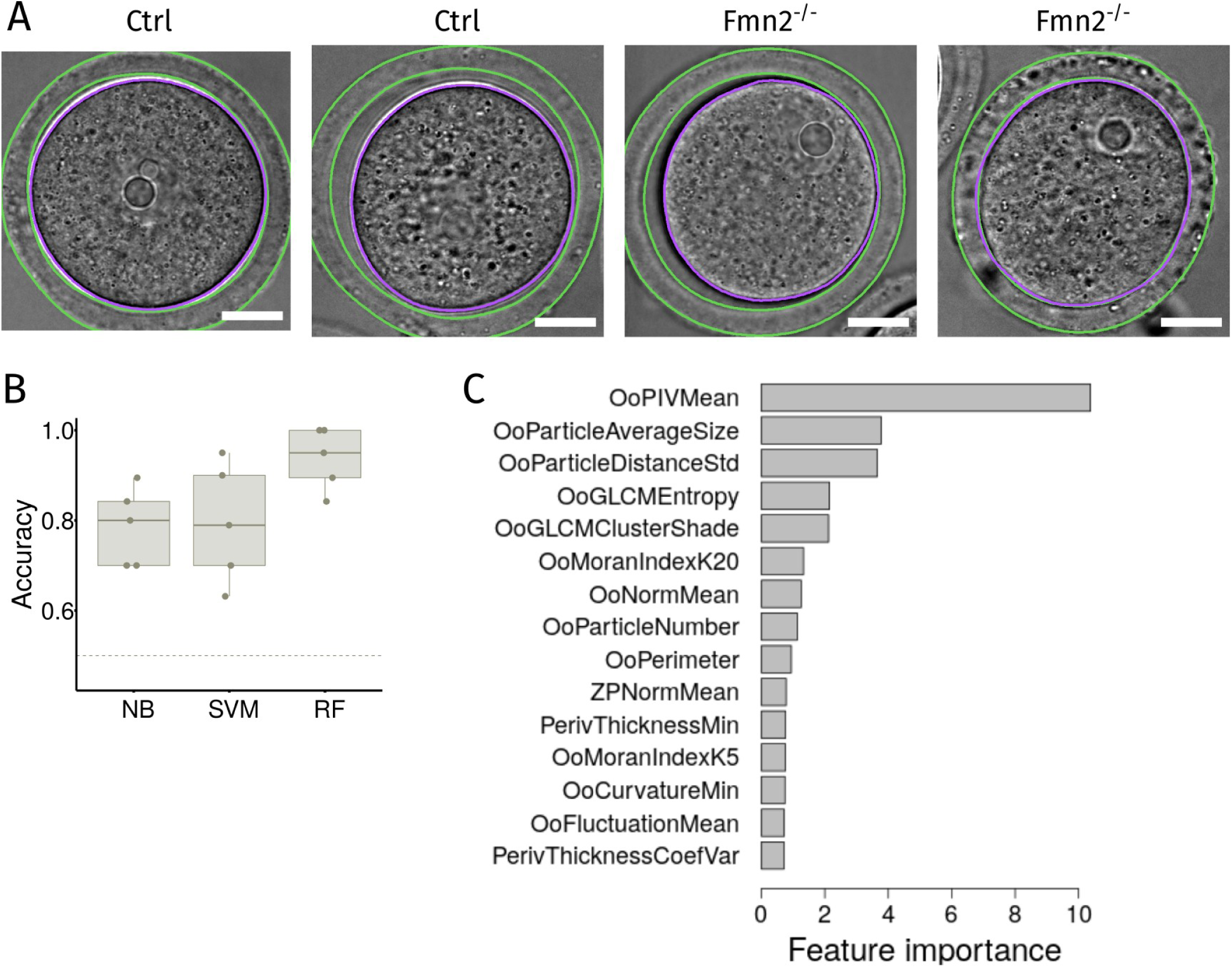
Phenotyping of Fmn2^-/-^ vs Control oocytes using our machine learning pipeline. A – Examples of segmentation of the oocyte contour (membrane, purple line and zona pellucida, green lines) obtained with our plugin on the new dataset. Scale bar is 20 µm. B – Comparison of several classification algorithms performance. 3 methods were tested in the analysis step of our pipeline: Naive Bayes (NB), Support Vector Machine (SVM) and Random Forest (RF). Their performance was measured by their Accuracy and compared with a 5-fold cross-validation technique. C – Features classified by their importance using our algorithm. The features were sorted by their importance calculated from the Gini indexes of each features in our Random Forest algorithm. We only display the scores of the 15 most important features.

**Figure S4:**
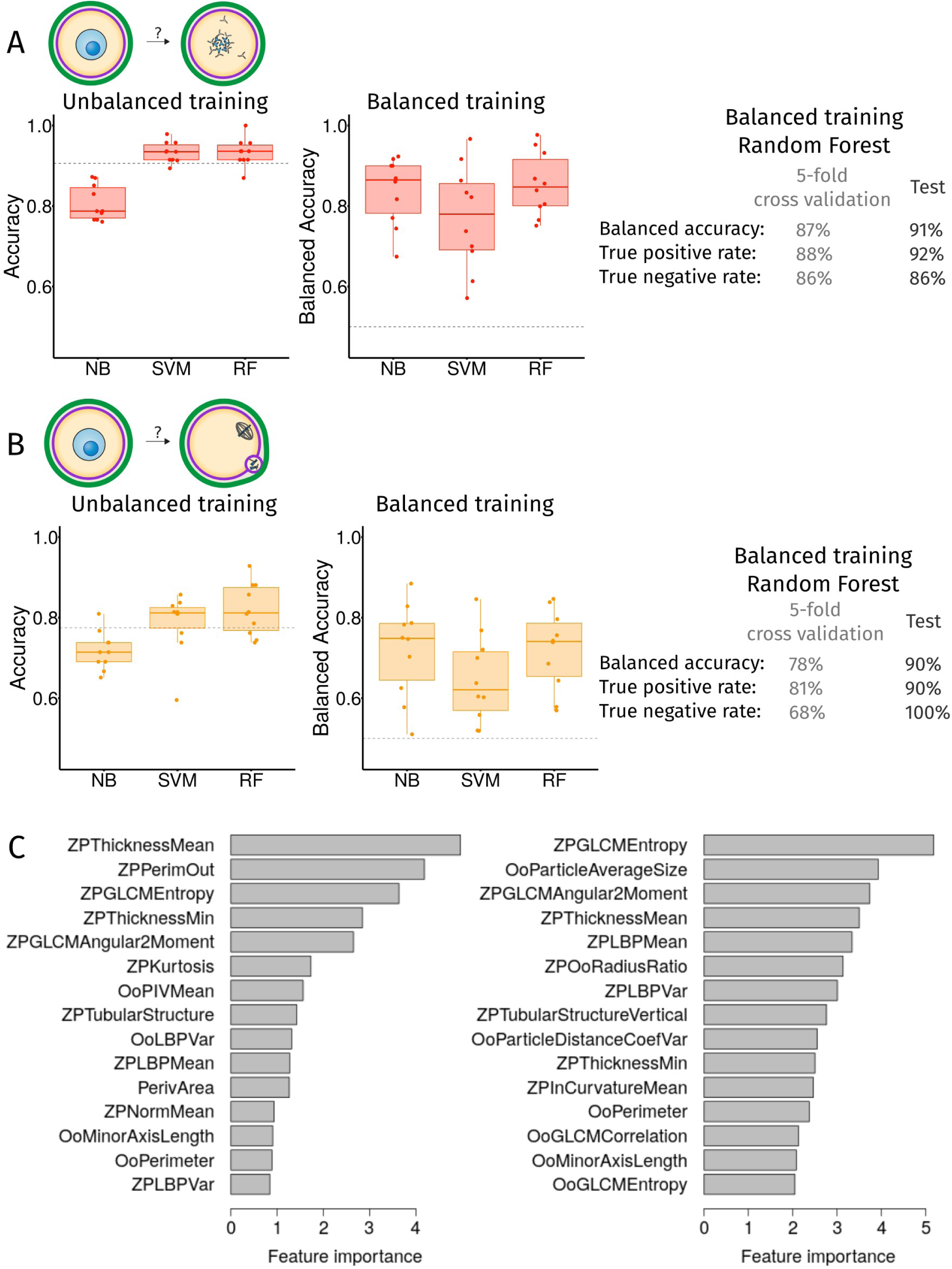
Prediction of NEBD failure and maturation defect. A – Scores of prediction of NEBD failure with 3 classification methods (Naive Bayes NB, Support Vector Machine SVM, Random Forest RF) assessed by 10-fold cross-validation. The training was done with the unbalanced dataset, thus considering classes frequency, and the score was consequently measured by the Accuracy (left graph). Dashed line represents the naive prediction score consisting in always predicting NEBD success. The training was also done with under-sampling the over-represented class (middle graph) and the score was measured by the balanced accuracy. The dashed line represents the naive prediction. Scores (balanced accuracy, true positive rate, true negative rate) obtained with the selected algorithm: Random Forest on balanced training for cross-validation and test on a new dataset. B - Scores of prediction of maturation defect with the same 3 classification methods (NB, SVM, RF) assessed by 10-fold cross-validation. The training was done with the unbalanced dataset, thus considering classes frequency, and the score was consequently measured by the Accuracy (left graph). Dashed line represents the naive prediction score consisting in always predicting maturation success. The training was also done with under-sampling the over-represented class (middle graph) and the score was measured by the balanced accuracy. The dashed line represents the naive prediction. Scores (balanced accuracy, true positive rate, true negative rate) obtained with the selected algorithm: Random Forest on balanced training for cross-validation and test on a new dataset. C - Features classified by their importance for NEBD failure (left panel) and maturation defect (right panel). Features sorted by importance calculated from the Gini indexes of each features in our Random Forest algorithm. We only display the scores of the 15 most important features. The significance of feature names are given in Supplemental File 1.

**Figure S5:**
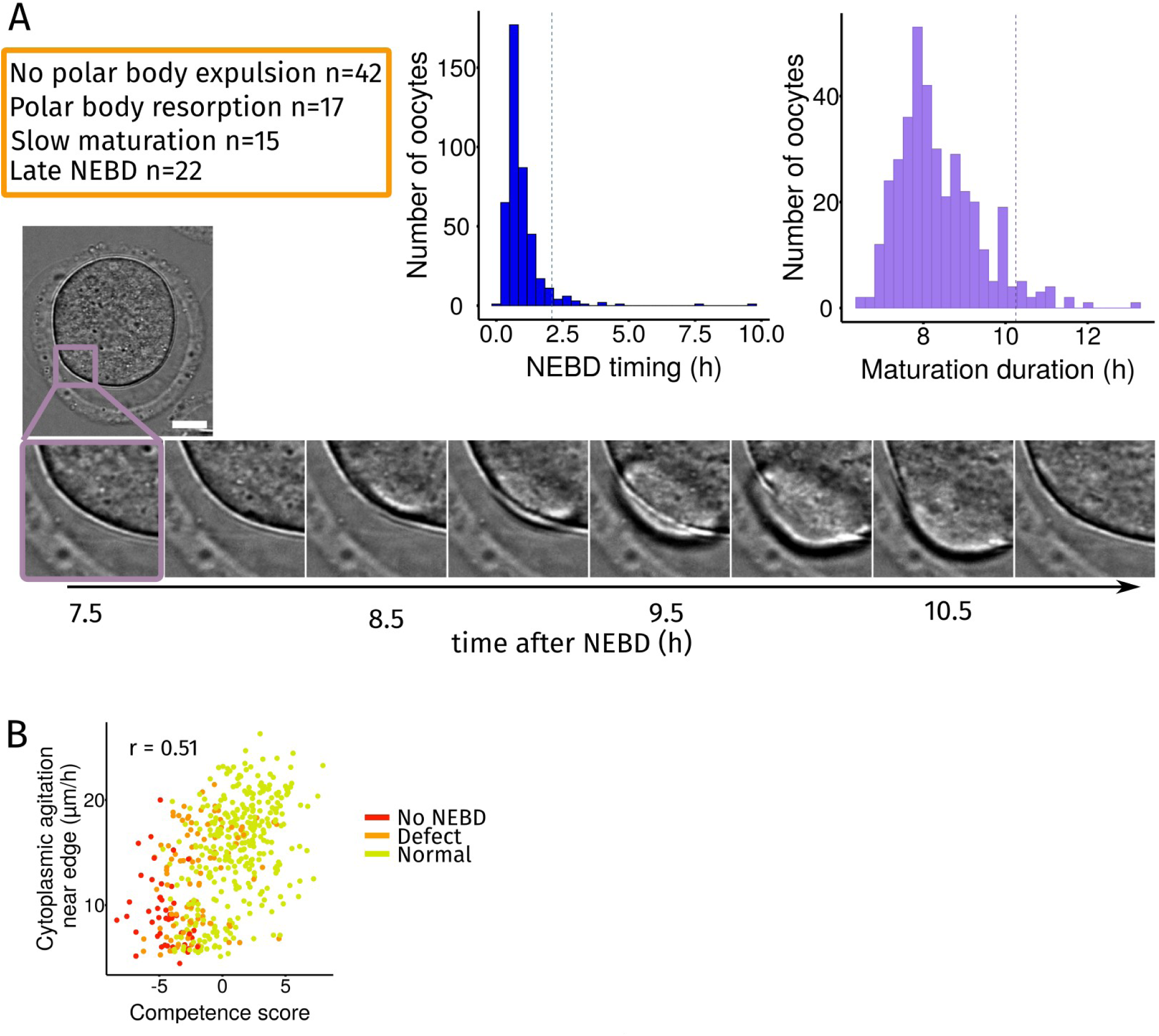
Characterization of oocyte maturation defects. A – Criteria used to classify a maturation as defective (top left panel, n is the number of oocytes for each defect). Histogram of NEBD timing in our dataset (middle panel). The dashed line represents the 95% quantile after which oocytes are considered as delayed in NEBD timing. Histogram of the time between NEBD and first polar body extrusion in our dataset (right panel). The dashed line represents the 95% quantile after which oocytes are considered to have a slow maturation. Time lapse of an example of polar body resorption (bottom panel). Scale bar is 20 µm. B – Correlation between cytoplasmic agitation and competence score. Cytoplasmic agitation in the area close to the edge of the oocyte (right panel) plotted against the oocyte competence score, for oocytes that do not enter into meiosis I (red, No NEBD), have a maturation defect (orange, Defect) or are normal (green, Normal). The Pearson correlation coefficients, r, are indicated in each graph with p-value < 10^-16 (Pearson’s product moment correlation coefficient) in both cases.

**Figure S6:**
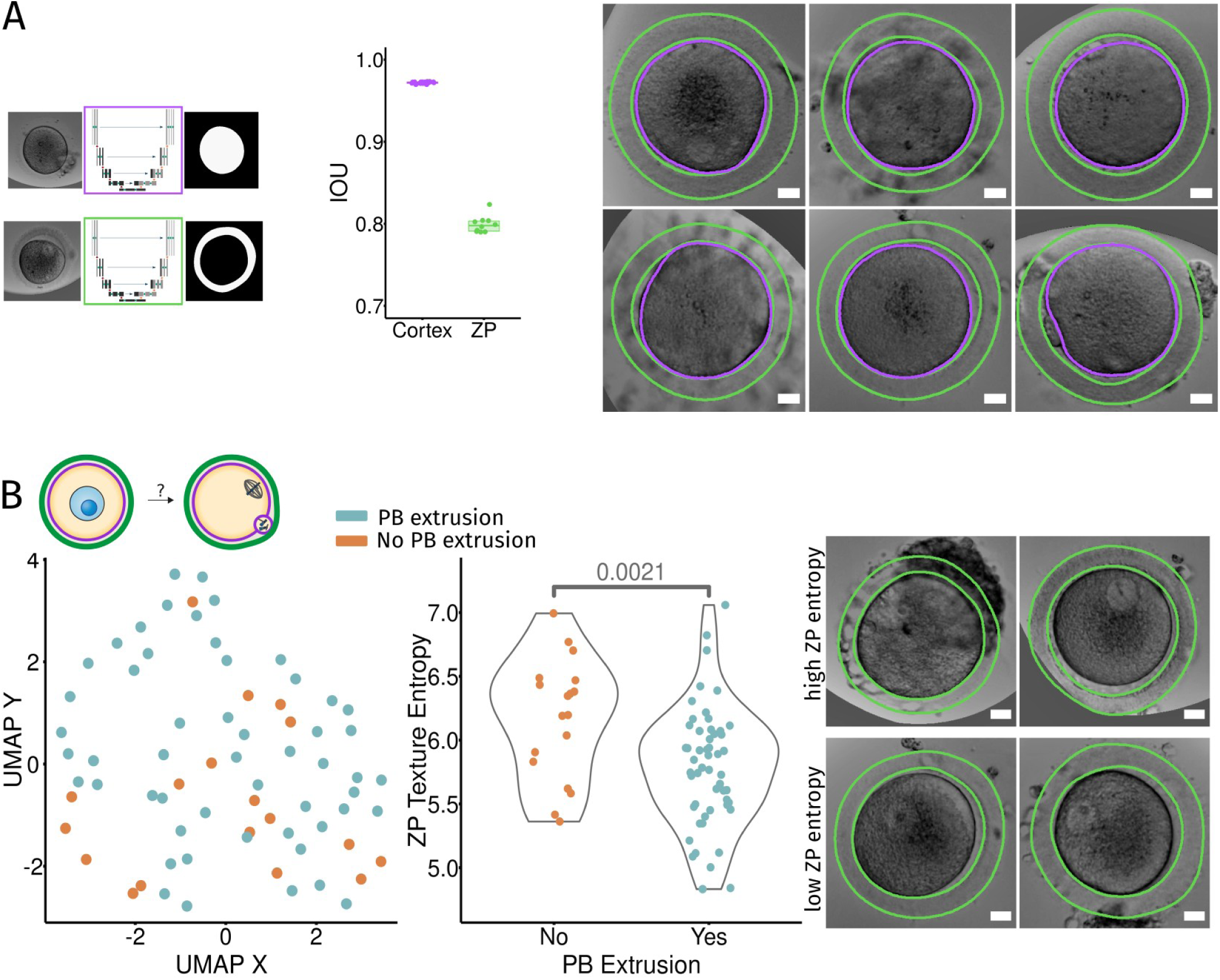
Characterization of human oocytes. A – Segmentation of human oocyte membrane (purple box) and zona pellucida (green box) using our trained neural network (left panel). Graph of the score of the segmentation measured as the Intersection Over Union (IOU) on the human test dataset (middle panel). Examples of segmentation of human oocytes (right panel). The oocyte membrane appears as a purple line and of the zona pellucida contours as green lines after segmentation. Scale bars are 20 µm. B – U-MAP projection of human oocyte features (left panel) for oocytes that will extrude a polar body (blue dots) and those who do not (orange dots). Values of the features at the beginning of the movies were used for all oocytes whether they had already entered meiosis or not. Comparison of zona pellucida texture entropy for oocytes that extrude a polar body or not (middle panel). Indicated p-value was calculated with a Kolmogorrov-Smirnov test. Examples of oocytes with high and low values for the entropy of the zona pellucida texture (right panel). Scale bars are 20 µm.

## Notes

### Competing Interest Statement

The authors have declared no competing interest.

